# Mutation-specific CAR T cells as precision therapy for IGLV3-21^R110^ expressing high-risk chronic lymphocytic leukemia

**DOI:** 10.1101/2023.09.29.560075

**Authors:** Florian Märkl, Christoph Schultheiß, Murtaza Ali, Shih-Shih Chen, Lukas Egli, Juliane Mietz, Obinna Chijioke, Lisa Paschold, Sebastijan Spajic, Anne Holtermann, Janina Dörr, Sophia Stock, Ignazio Piseddu, David Anz, Marcus Dühren-von Minden, Tianjiao Zhang, Thomas Nerreter, Michael Hudecek, Nicholas Chiorazzi, Sebastian Kobold, Mascha Binder

## Abstract

The concept of precision cell therapy targeting tumor-specific mutations is appealing but requires surface-exposed neoepitopes, which is a rarity in cancer. B cell receptors (BCR) of mature lymphoid malignancies are exceptional in that they harbor tumor-specific-stereotyped sequences in the form of point mutations that drive self-engagement of the BCR and autologous signaling. Here, we used a BCR light chain neoepitope defined by a characteristic point mutation (IGLV3-21^R110^) for selective targeting of a poor-risk subset of chronic lymphocytic leukemia (CLL) with chimeric antigen receptor (CAR) T cells. We developed murine and humanized CAR constructs expressed in T cells from healthy donors and CLL patients that eradicated IGLV3-21^R110^ expressing cell lines and primary CLL cells, but not polyclonal healthy B cells. In vivo experiments confirmed epitope-selective cytolysis in xenograft models using engrafted IGLV3-21^R110^ expressing cell lines or primary CLL cells. We further demonstrate in two humanized mouse models lack of cytotoxicity towards human B cells. These data provide the basis for novel avenues of resistance-preventive and biomarker-guided cellular targeting of functionally relevant lymphoma driver mutations sparing normal B cells.

## Introduction

Chronic lymphocytic leukemia (CLL) is a paradigmatic low-grade lymphoma in which the B cell receptor (BCR) plays a central biological role. The BCR landscape of CLL has been extensively studied both immunogenetically and functionally. These studies revealed recognition of distinct self-antigens through stereotyped complementarity-determining region 3 (CDR3) sequence motifs, oligomeric membrane organization as well as autonomous signaling through BCR-BCR interactions (e.g.,^1–5^). Some patients from stereotypic BCR subsets are poor-risk with only limited long-term clinical benefit with established approaches including those that target the BCR pathway (e.g.,^6^).

Advanced immunotherapeutic approaches such as chimeric antigen receptor (CAR) modified T cells are under clinical investigation for patients with poor-risk CLL. CAR T cells against CD19, a component of the BCR complex, can provide significant activity in patients suffering from advanced or refractory CLL but the rate of complete remissions and long-term responses remain well behind that observed in other lymphoma types^7–13^. Suboptimal outcomes are due to CD19 antigen loss, CAR T cell loss or dysfunction, and complete eradication of the B cell lineage that causes clinically relevant immunosuppression in these studies. Along these lines, it appears that CAR T cells can in principle be effective in CLL. However, an ideal target should be tumor-specific and of high functional relevance to prevent downregulation or loss under selective pressure. Such novel precision approaches may help to achieve durable benefit eventually even resulting in cure, as seen in other indications.

The discovery of a landscape of disease-specific sequence motifs in BCRs expressed by the malignant CLL clone opened new avenues for targeted cell therapy that may eventually be translated to other types of lymphomas. Here, we provide first proof-of-concept for the activity of bona-fide tumor-specific CAR T cells for high-risk patients with CLL that express the IGLV3-21^R110^ BCR light chain. The IGLV3-21^R110^ subset typically shows an aggressive clinical course^6^. IGLV3-21^R110^ is expressed in 10-15% of unselected CLL patients, but overrepresented in treatment-requiring CLL. Functionally, the G-to-R exchange at position 110 of the IGLV3-21 light chain – along with several conserved amino acids also in the heavy chain – confers autonomous signaling capacity to the BCR by mediating self-interactions^14–21^. Since the IGLV3-21^R110^ BCR is CLL-specific and represents a critical tumor driver, we reasoned that targeting this receptor would spare normal B cells and may have a low risk of epitope escape. At the same time, the lack of persistent B cell aplasia may be of advantage in terms of infection-mediated complications and preserved responses to vaccination. We developed IGLV3-21^R110-^targeted CAR T cells including humanized variants thereof and demonstrate in vitro and in vivo, including against primary CLL samples, selective targeting and eradication, leaving the healthy B cell compartment untouched. These results underpin the potential value of such precision approach and warrant clinical investigations.

## Materials and Methods

### Patient cohort

Blood samples from 158 CLL patients were collected after informed consent as approved by the ethics committees of the Universities of Hamburg–Eppendorf, Freiburg and Halle-Wittenberg. Peripheral mononuclear cells (PBMCs) were isolated by Ficoll gradient centrifugation, resuspended in FCS + 10% DMSO and cryopreserved in liquid nitrogen. IGLV3-21^R110^ expressing CLL was characterized by flow cytometry and next-generation sequencing (NGS) of the light chain loci as described below^22–31^.

### IGLV3-21^R110^ Flow Cytometry

IGLV3-21^R110^ expression was tested with an APC-labelled IGLV3-21^R110^-specific antibody (AVA Lifescience GmbH, Denzlingen, Germany). Twenty cases were additionally analyzed with the ApLife^TM^ FastScreen_CLL_ assay that uses CD19, CD5 and IGLV3-21^R110^ antibodies in addition to labelled spheres to define cut-off levels comparable throughout different measurements.

### IGLV3-21^R110^ Next-generation sequencing (NGS)

IGL repertoires were profiled as described^22,25–28,30^. The IGL primer pool was adapted from^32^ to cover the complete IGLJ (FR4) region including the first nucleotide of the triplet for amino acid position 110 at the junction of IGLJ and IGLC. The sequences of the new reverse primers are (5’–3’): GTGAGACAGGCTGGG, CAAGAGCGGGGAAGG, CAACTTGGCAGGGAAAG, GGGAGACCAGGGAAG, TCACCCTAGACCCAAAAG. The MiXCR framework^33^ with the IMGT library^34^ as reference for sequence alignment was used for clonotype assembly. Amino acid position 110 was defined by nucleotide 28 of the FR4 region.

### Cell lines and primary CLL and healthy donor blood cells

OCI-Ly1 and NALM-6 were purchased from the DSMZ (German Collection of Microorganisms and Cell Cultures GmbH). Luciferase (Luc) overexpressing cell line NALM-6 Luc was previously described^35^. OCI-Ly1 Luc-GFP was generated as described^36^. For ectopic expression of the IGLV3-21^R110^ light chain, the coding sequence was cloned into the Lentiviral Gene Ontology (LeGO) vector LeGO-iC2-Puro+ via AsiSI/EcoRI (for expression in OCI-Ly1) and the retroviral vector pMP71 (for expression in NALM-6 Luc), respectively^37^.

### CAR constructs

The CAR construct derived from the murine single-chain variable fragment (scFv) of the IGLV3-21^R110^-specific antibody from AVA Lifescience GmbH (Denzlingen, Germany). It was cloned into the retroviral vector pMP71 containing CD28 and CD3ζ costimulatory domains. The scFv sequence derived from the murine anti-IGLV3-21^R110^ antibody was humanized and cloned via *Nhe*I and *Rsr*II into an established lentiviral CD19 CAR vector containing 4-1BB and CD3ζ costimulatory domains^38^. Human thyrotropin receptor-directed CAR T cells (αTSHR-CAR)^39^ as well as a published CD19-targeting CAR (αCD19-CAR)^38^ served as control. These CARs contained a truncated epidermal growth factor receptor (EGFRt) for sorting of CAR-expressing cells^40^. All sequences are listed in Supplementary Table S1.

### Virus production

Lentivirus production was performed as described earlier^41^. For retrovirus production, 293Vec-Galv and 293Vec-RD114 cell lines^42^ were used (kind gift of Manuel Caruso, Québec, Canada). Retroviral pMP71 vectors (kindly provided by C. Baum, Hannover) carrying the sequence of the relevant receptor were stably introduced in packaging cell lines^36^. Single cell clones were generated and indirectly screened for virus production by determining transduction efficiency of primary T cells. This method was used to generate the producer cell lines 293Vec-RD114 for scFv-R110-CD28-CD3ζ (αR110-mCAR), EGFRvIII-CD28-CD3ζ (E3 synthetic agonistic receptor (E3-SAR)) and scFv-CD19-CD28-CD3ζ (αCD19-mCAR (WO2015187528A1)).

### Generation of CAR-expressing primary human T cells

Pan T cells were isolated from healthy donor or CLL patient-derived whole blood (Pan T Cell Isolation Kit and Auto MACS Quant, Miltenyi, Bergisch Gladbach, Germany), stimulated with CD3/CD28 T-cell activation Dynabeads (Life Technologies, Carlsbad, USA) at a 1:1 bead to cell ratio, and lentivirally transduced 24 hours later at a multiplicity of infection of 1.5 or retrovirally transduced 48 hours after isolation^43^. All T cells were expanded in complete T cell medium supplemented with penicillin–streptomycin (100 U/mL; Life Technologies, Carlsbad, USA)) and fed IL-2 (50 U/mL; Stem Cell Technologies, Vancouver, Canada) every 48 hours. Dynabeads were removed day 6 after isolation.

### Antibody and scFv affinity ranking

The affinity of the murine anti-IGLV3-21^R110^ antibody from AVA Lifescience and the humanized single-chain variable fragment (scFv) was determined using flow cytometry and the TKO cell model^44^. For ectopic expression of the IGLV3-21^R110^ light chain in TKO cells, the coding sequence was cloned into the vector pMIZYN. Transduction was performed after generation of lentiviral particles in 293T cells as described above. For binding, 2 x 10^5^ TKO cells were seeded as duplicates in 96-well plates and incubated with serial dilutions of antibody/scFv for 30 min at 4°C followed by secondary detection (anti-human-IgG1-APC, Clone IS11-12E4.23.20, Miltenyi, Bergisch Gladbach, Germany) and quantification on a MACSQuant Analyzer 10 flow cytometer (Miltenyi, Bergisch Gladbach, Germany).

### In vitro cytotoxicity assay and cytokine quantification

For Incucyte S3 assays, target cells seeded at 2 × 10^4^ cells/well in a 96-well plate were co-incubated with effector cells at effector-to-target (E:T) ratio 5:1 in complete media. Primary CLL target cells were isolated by Ficoll gradient centrifugation. Polyclonal control B cells from healthy donors were isolated by Dynabeads™ CD19 isolation kit from Invitrogen after Ficoll gradient centrifugation. CAR T cell-mediated tumor cell cytotoxicity was assessed using the Incucyte Caspase-3/7 Reagent (BioScience, Essen, Germany).

Other cytotoxicity assays were performed using a flow cytometry-based readout (BD LSRFortessa (BD Biosciences, New Jersey, USA)) after 48 hours of coculture with human CAR T cells. Dead cells were stained using the violet fixable viability dye (BioLegend, San Diego, USA) for 10 minutes at room temperature. Following this, cell surface proteins were stained for 20 minutes at 4 °C. Tumor cells were quantified by using an anti-CD19 antibody (6D5). For the quantification of the CAR T cells, antibodies against CD4 (OKT4), CD8a (RPA-T8), EGFR (A-13) (all from BioLegend, San Diego, USA) and c-Myc (SH1-26E7.1.3, Miltenyi Biotech, Bergisch-Gladbach, Germany) were used. Furthermore, luciferase-based toxicity assays were performed using Bio-Glo Luciferase Assay System (Promega Corporation, USA).

In addition, cytokine measurements were done by ELISA (BD Biosciences, Frankllin Lakes, USA) or a bead-based immunoassay technology (LEGENDplex, Biolegend, San Diego, USA). Values below the limit of detection were considered zero.

### CAR T cell in-vivo assays

Experiments were performed with female NSG (NOD.Cg-Prkdc^scid^ Il2rg^tm1WjI^/SzJ) mice aged 2-3 months from Janvier Labs (n=60) according to the regulations of the Regierung von Oberbayern. NALM-6 Luc-R110 xenograft models were established in NSG mice following the intravenous (i.v.) injection of 10^5^ tumor cells in 100 μL PBS. Animals were randomized into treatment groups according to tumor burden. Experiments were performed by a scientist blinded to treatment allocation and with adequate controls. No time points or mice were excluded from the experiments presented in the study. For adoptive cell therapy (ACT) studies, 4-5 x 10^6^ active CAR T cells were injected i.v. in 100 µl PBS. Tumor burden was measured using a luciferase-based IVIS Lumina X5 imaging system (PerkinElmer, Waltham, USA). Survival analyses were recorded in Kaplan Meyer plots.

### Patient-derived CLL xenograft

T cells derived from PBMCs of patient CLL425 were activated with CD3/CD28 dynabeads and IL-2 for 3 days. The activated T cells were then mixed with CLL425 PBMCs at the ratio of 1 to 40. Total 0.5 x 10^6^ T cells and 20 x 10^6^ PBMCs were intravenously injected into each NSG (NOD/SCID/IL2rγnull) mouse, and a total of 12 mice were injected. 10 days after, mice were randomly split into two groups, 6 mice were intraperitoneally injected with 7 million CLL-αR110-CAR T cells (n = 6), or untransduced T cells (UTD) generated from the same patient. Three weeks later, all the mice were sacrificed, blood, spleen and bone marrow samples were collected for the number of T and CLL cells. Animal studies were performed in accordance with experimental protocols approved by the Institutional Animal Care and Use Committee (IACUC) of the Feinstein Institute for Medical Research. Single cell suspension prepared from these tissues were then stained and analyzed by flow cytometry for the number of human CD45^+^CD19^+^CD5^+^ CLL B cells and CD45^+^CD19^-^CD5^+^ T cells.

### Humanized mouse models

Human PBMCs were isolated by gradient centrifugation. Female NSG mice were purchased from Janvier Labs (in total: n = 25) and injected at 6-8 weeks i.v. with 20 x 10^6^ PBMCs. On day seven the mice were injected i.v. with 1.5 x 10^6^ active αR110-CAR T or αCD19-CAR T cells. The frequency of CD19^+^ CD20^+^ cells of all CD45^+^ cells was analyzed by flow cytometry (d8,13). In a separate assay, human PBMCs were labelled with the fluorescent membrane marker PKH26 (Sigma-Aldrich, MINI26-1KT). For the in vivo killing assay, 3-5 month old NFA2 (NOD.Cg-*Rag1^tm1Mom^ Flt3^tm1Irl^ Mcph1^Tg(HLA-A2.1)1Enge^ Il2rg^tm1Wjl^*/J) mice were injected i.p. with 2.5 x 10^6^ PKH26-labelled PBMC and 1.25 x 10^6^ αR110-CAR T or αCD19-CAR T cells (in total n = 18). Mice were sacrificed after 16 hours, peritoneal cells were harvested by peritoneal lavage and quantified using flow cytometry. Mice in which either PBMC or CAR T cells could not be detected by flow cytometry were excluded from the dataset to control for unsuccessful i.p. injections. The following markers were used: anti-human CD45-Pacific Blue (Biolegend, 368540), anti-mouse CD45 (30F11, Biolegend, San Diego, USA), anti-human CD19 (HIB19, Biolegend, San Diego, USA), anti-human CD3 (UCHT1, Biolegend, San Diego, USA), anti-human CD14 (TuK4, Invitrogen, Carlsbad, USA), anti-human CD56 (NCAM16.2, BD Franklin Lakes, USA), Zombie Aqua live/dead stain (423102, Biolegend, San Diego, USA).

### Statistical analysis

One-tailed student’s t-test was used for comparisons between two groups. A log-rank (MantellZICox) test was used to compare survival curves. All statistical tests were performed with GraphPad Prism software (v8.3.0). P values are represented as * < 0.05, ** < 0.01, *** < 0.001 and **** < 0.0001. No statistical methods were used to predetermine sample size.

## Results

### Anti-IGLV3-21^R110^ CAR T cells exhibit epitope-selective tumor cell lysis in vitro

To target the CLL-specific IGLV3-21^R110^ light chain mutation, we first utilized a murine IGLV3-21^R110^–specific antibody to generate a 2^nd^ generation CAR with CD28-CD3ζ signaling domain for retroviral transduction of primary human healthy donor (HD) T cells (HD-αR110-mCAR T cells) (Figure 1A, 2A;. supplemental Figure 1A). For proof-of-principle experiments, we transduced Luciferase (Luc) overexpressing NALM-6 cells (NALM-6 Luc)^35^ with the pMP71 retroviral vector encoding IGLV3-21^R110^ to generate a target cell line with constitutive surface expression of a hybrid BCR containing the IGLV3-21^R110^ light chain (NALM-6 Luc-R110). Co-culture of NALM-6 Luc-R110 cells with HD-αR110-mCAR T cells showed epitope-selective lysis of IGLV3-21^R110^-expressing lymphoid target cells, while control NALM-6 Luc cells were unaffected (Figure 1B). CD19-directed CAR T cells (HD-αCD19-mCAR) equally lysed both cell lines (Figure 1B), while co-incubation with an unrelated CAR product (HD-E3-SAR ctrl)^45,46^ or untransduced primary human T cells (UTD) failed to lyse NALM-6 lines (Figure 1B). These specific lysis patterns were paralleled by equivalent IFN-γ secretion patterns (Figure 1C). Notably, HD-αR110-mCAR T cells preferentially expanded in co-culture with NALM-6 Luc-R110 cells (supplemental Figure 1B), indicating good functionality of HD-αR110-mCAR T cells. Importantly, HD-αR110-mCAR T cells were not activated by and did not lyse polyclonal human B cells (supplemental Figure 1C).

**Figure 1.**
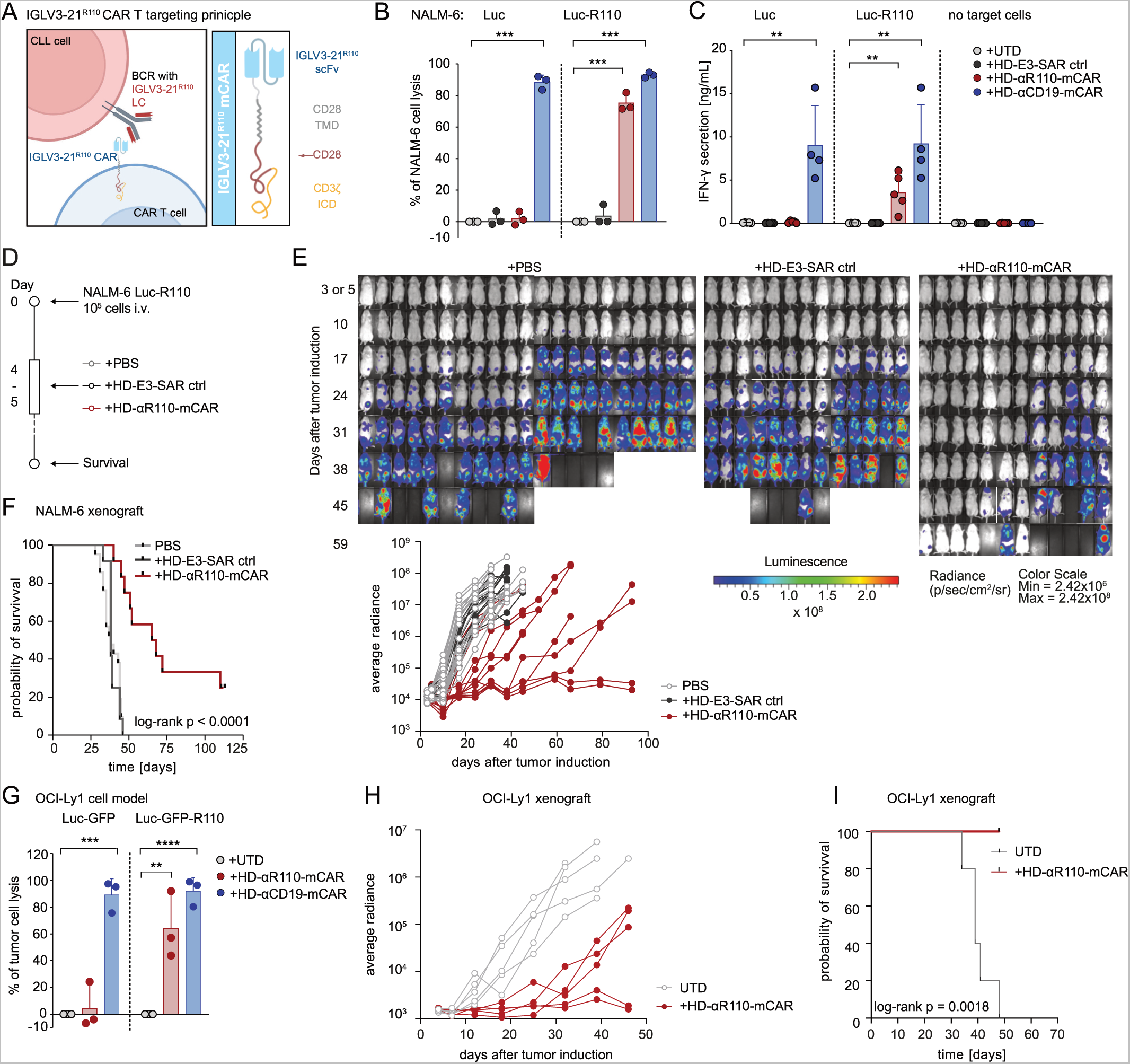
Development of a chimeric antigen receptor (CAR) T cell targeting principle against the IGLV3-21^R110^ neoepitope. (A) Schematic representation of the IGLV3-21^R110^ CAR T targeting principle and the murine CAR construct. BCR, B cell receptor. LC, light chain. TMD, transmembrane domain. ICD, intracellular domain. (B) Percent tumor cell lysis based on a bioluminescence readout after 48h co-culture of NALM-6 Luc or IGLV3-21^R110^ expressing NALM-6 Luc-R110 cells with indicated CAR T cells or untransduced T cells (UTDs) in an effector to target (E:T) ratio of 0.1:1. (C) Quantification of IFN-γ secretion in cell culture supernatants after 48h co-culture of NALM-6 Luc model. (D) Schematic representation of the workflow for the NALM-6 Luc-R110 xenograft mouse model. (E) Bioluminescence measurements of NSG mice to assess *in vivo* activity of HD-αR110-mCAR T cells from healthy donors. NALM-6 Luc-R110 growth curves are shown based on *in vivo* bioluminescence imaging (days 3 or 5, 10, 17, 24, 31, 38, 45, 52, 59, 65 or 66, 79. 93). NSG mice were injected i.v. with NALM-6 Luc-R110 cells and treated either four or five days later with HD-αR110-mCAR T cells (n = 12), control HD-E3-SAR ctrl T cells (n = 12) or PBS vehicle solution (n = 21). (F) Kaplan-Meier survival plot of NALM-6 Luc-R110 mouse model. (G) Percent tumor cell lysis based on a bioluminescence readout after 48h co-culture of OCI-Ly1 Luc-GFP or IGLV3-21^R110^ expressing OCI-Ly1 Luc-GFP-R110 cells with indicated CAR T cells in an effector to target (E:T) ratio of 0.1:1. (H) Bioluminescence measurements of NSG mice engrafted with OCI-Ly1 Luc-GFP-R110 cells to assess *in vivo* activity of HD-αR110-mCAR T cells from healthy donors. OCI-Ly1 Luc-GFP-R110 growth curves are shown based on *in vivo* bioluminescence imaging (days 4, 7, 12, 18, 25, 32, 39, 46). NSG mice were injected i.v. with OCI-Ly1 Luc-GFP-R110 cells and treated either eight days later with HD-αR110-mCAR T cells or UTD (for all n = 5). (I) Kaplan-Meier survival plot of OCI-Ly1 Luc-GFP-R110 mouse model. All bar plots represent the indicated mean ± SD calculated from three independent experiments. Statistics: one-sided t test. For statistical analysis of survival data, the log-rank test was applied.

**Figure 2.**
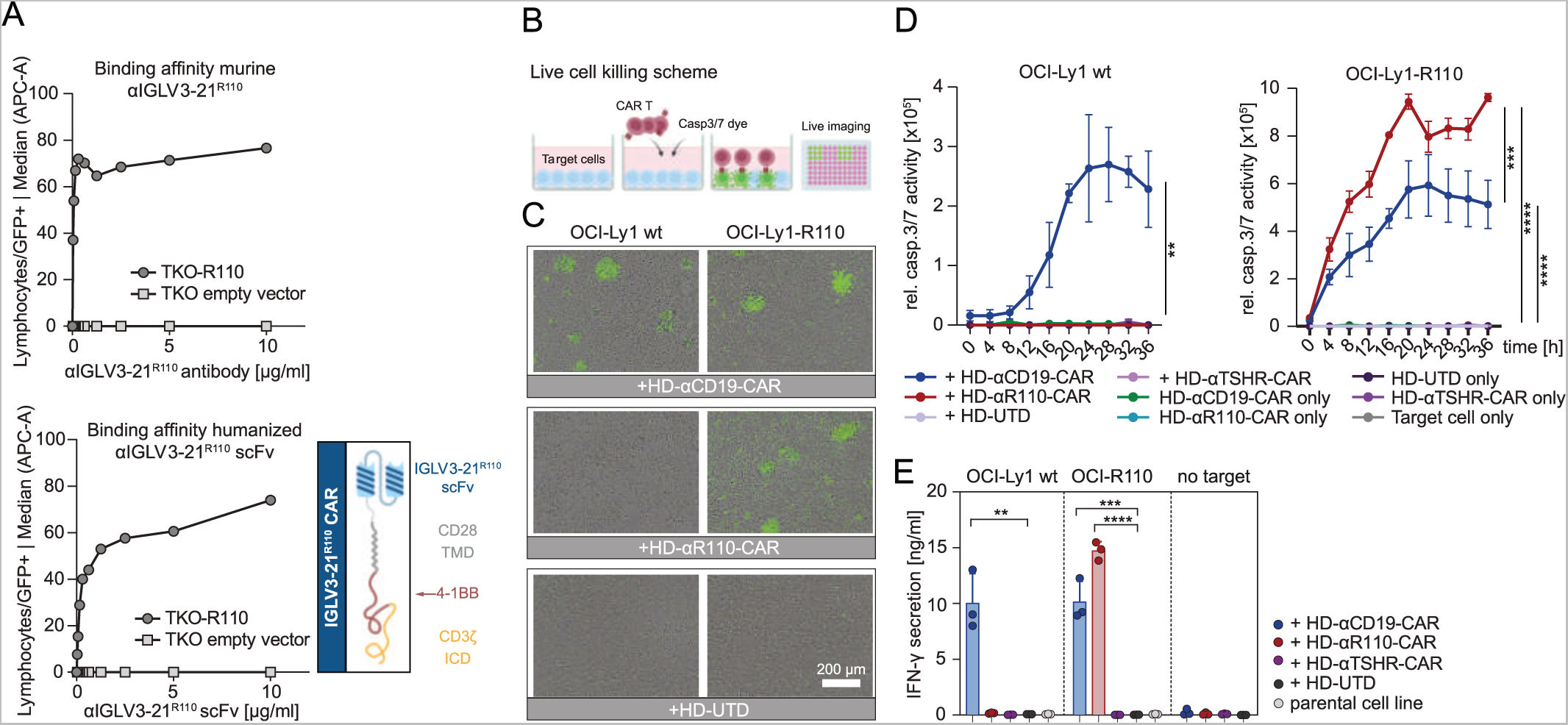
Humanization of the binding moiety of chimeric antigen receptor (CAR) T cells against the IGLV3-21^R110^ neoepitope. (A) Determination of median binding affinities of the murine anti-IGLV3-21^R110^ antibody or the corresponding humanized scFv construct. Serial dilutions were incubated with TKO cells expressing the IGLV3-21^R110^ antigen and binding intensity was quantified using flow cytometry. (B) Schematic representation of the applied live cell killing assay using the Incucyte system and the Caspase-3/7 fluorescence dye. (C)-(D) Cytolysis of IGLV3-21^R110^ expressing OCI-Ly1-R110 cells and OCI-Ly1 wild-type (wt) cells by healthy donor anti-IGLV3-21^R110^ CAR T cells with a humanized scFv sequence (HD-αR110-CAR) with an E:T ratio of 5:1. Cytolysis was monitored over time as caspase 3/7 activity (relative intensity of green fluorescence) in co-cultures using Incucyte S3. Anti-thyroid stimulating hormone receptor (TSHR) CAR T cells (HD-αTSHR-CAR) and untransduced T cells (UTD) served as controls. All conditions indicated have been plotted, negative control conditions overlap. (C) Representative images show CAR T cell-mediated cytolysis after 24h. (E) Quantification of IFN-γ in co-culture supernatants after 24h incubation of indicated target/effector cell combinations. All bar plots represent the indicated mean ± SD. Statistics: one-sided t test.

### αR110-mCAR CAR T cells are efficacious in xenograft R110 -models

We next tested the efficacy of HD-αR110-mCAR T cells in NSG mice engrafted with NALM-6 Luc-R110 cells (Figure 1D). Bioluminescence imaging showed substantial reduction of NALM-6 Luc-R110 outgrowth in mice treated with HD-αR110-mCAR T cells (Figure 1E), accompanied by prolonged survival and disease eradication in 17% of treated mice (Figure 1E, F). We next set up a second xenograft model using the same engineering strategy to generate OCI-Ly1 lymphoma cells expressing the IGLV3-21^R110^ light chain (OCI-Ly1 Luc-GFP-R110). We also included CD19-directed CAR T cells (HD-αCD19-mCAR T cells) for head-to-head comparisons. As observed for the NALM-6 model, HD-αR110-mCAR T cells selectively lysed OCI-Ly1 Luc-GFP-R110 cells in co-culture experiments, while control OCI-Ly1 Luc-GFP cells were unaffected (Figure 1G). HD-αCD19-mCAR T cells lysed OCI-Ly1 Luc-GFP cells independently of IGLV3-21^R110^ light chain status (Figure 1G). Mice engrafted with OCI-Ly1 Luc-GFP-R110 cells controlled and to some extent even cleared disease, when injected with HD-αR110-mCAR T (Figure 1H). Importantly, these mice survived the 50 days of the experiment in both settings, indicative of comparable activity (Figure 1I).

### Humanization of the anti-IGLV3-21^R110^ scFv sequences preserves functionality

Given the potential immunogenicity of xenogeneic protein-components such as a murine scFv^47^ we next humanized the anti-IGLV3-21^R110^ scFv sequence to generate a CAR construct with potentially lower immunogenicity (Figure 2A; supplemental Table 1). We used a flow cytometry based affinity ranking assay and IGLV3-21^R110^-expressing TKO cells^44^ to compare the concentration-dependent binding capabilities of the murine anti-IGLV3-21^R110^ antibody and the humanized scFv. As shown in Figure 2A, humanization did not compromise the binding affinity of the purified scFv fragment. Next, we cloned the humanized scFv fragment in a 2^nd^ generation CAR backbone with 4-1BB-CD3ζ costimulatory domain (Figure 2A) and lentivirally transduced this construct into human T cells for CAR T generation (HD-αR110-CAR) (supplemental Figure 1D). To test their functionality in vitro, we performed co-culture killing assays using OCI-Ly1 cells overexpressing IGLV3-21^R110^ (OCI-Ly1-R110) and a fluorescent reporter dye indicating caspase3/7-mediated lymphoma cell apoptosis (Figure 2B). The obtained live cell imaging data suggested that HD-αR110-CAR T cells selectively target OCI-Ly1-R110 cells, while HD-αCD19-CAR lyse OCI-Ly1 cells independently of IGLV3-21^R110^ status (Figure 2D). Co-culture with an unrelated CAR targeting the human thyroid stimulating hormone receptor (TSHR) or untransduced primary T cells did not affect OCI-Ly1 viability (Figure 2B). Selectivity of the HD-αR110-CAR T cells was also suggested by IFN-γ-release patterns (Figure 2D-E).

### Healthy donor or CLL patient-derived anti-IGLV3-21^R110^ CAR T cells efficiently target primary CLL cells

To come closer to patient settings, we next asked if this targeting principle is also applicable to primary CLL cells. To identify eligible individuals for target cell isolation, we first screened a cohort of 158 CLL patients for IGLV3-21 status. Using light chain NGS and flow cytometry^31^ we identified 17 IGLV3-21^R110^ cases (Figure 3A). Since light chain sequencing is not a clinical standard in CLL and staining results with the murine IGLV3-21^R110^ antibody under varying conditions of routine flow cytometry labs may differ, we set up a more standardized synthetical particle-based flow cytometry IGLV3-21^R110^ detection method. Twenty samples from the above mentioned CLL cohort were randomly selected for quantification using normalization with synthetical beads, which showed 100% concordance with prior conventional typing results (Figure 3A). Next, we selected two treatment-naïve CLL patients with or without IGLV3-21^R110^ mutation for target cell isolation. Co-culture with HD-αR110-CAR T cells showed selective lysis of IGLV3-21^R110^-positive CLL cells after 24 hours as previously demonstrated with neoepitope-transduced cell lines (Figure 3B). HD-αCD19-CAR T cells lysed CLL cells from all included patients (Figure 3B). Co-culture with HD-αTSHR-CAR T cells or untransduced T cells had no effect on the co- culture (Figure 3B). Cytolysis of primary CLL cells was paralleled by IFN-γ release (Figure 3C). We then generated CAR T cells from primary T cells of two patients with CLL – one patient with active CLL (CLL433) and one in remission (CLL453) – to demonstrate their cytotoxic capacity despite the known dysfunction of T cells in this disease. Patient-derived CLL-αR110-CAR T cells showed selective lysis of OCI-Ly1-R110, while CLL-αCD19-CAR T cells from the same patients exhibited cytolysis irrespectively of the neoepitope (Figure 3D). Next, we tested efficacy of CLL-αR110-CAR T cells derived from patient CLL425 with IGLV3-21^R110^-positive active CLL in an autologous setting using a primary CLL xenograft mouse model (Figure 3E). Application of autologous CLL-αR110-CAR T cells reduced primary CLL but not T cell load at the three week end-point in spleen and bone marrow (Figure 3F).

**Figure 3.**
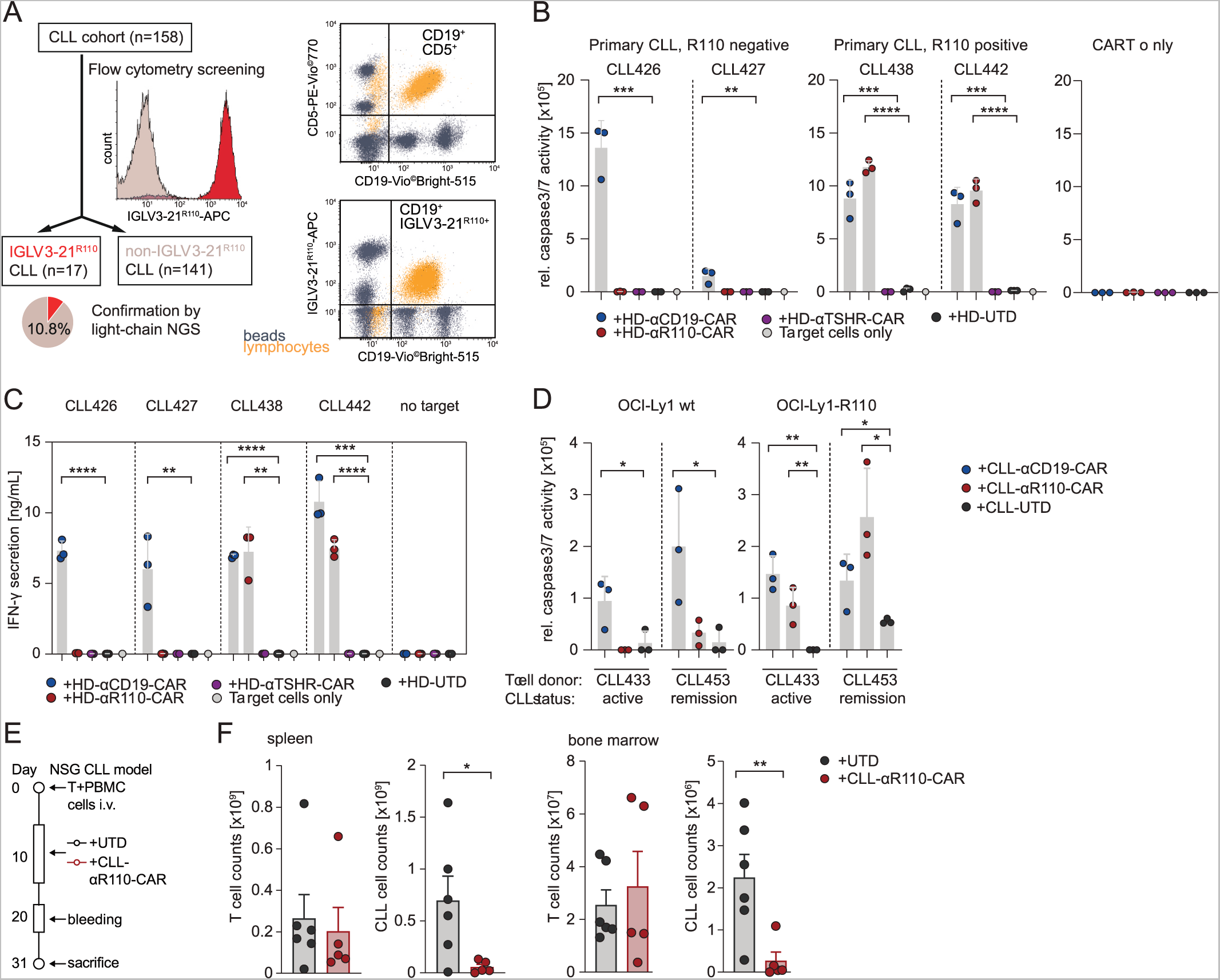
In-vitro and in-vivo activity of IGLV3-21^R110^ healthy donor and CLL patient-derived CAR T cells against primary chronic lymphocytic leukemia (CLL) cells. (A) Screening work-flow of 158 CLL patients for the IGLV3-21^R110^ light chain including a standardized bead-based assay. Exemplary results of one IGLV3-21^R110^-positive and one IGLV3-21^R110^-negative CLL case in a single color flow cytometric assay using APC-labelled IGLV3-21^R110^-specific antibody shown as histogram. Exemplary staining of one IGLV3-21^R110^-positive CLL case with a bead-based assay (ApLife^TM^ FastScreen_CLL_) with triple staining of CD19, CD5 (CD19^-^CD5^+^ unaffected T cells) and IGLV3-21^R110^. (B) Cytolysis of freshly isolated primary CLL cells from IGLV3-21^R110^-positive (CLL438, CLL442) and IGLV3-21^R110^-negative (CLL426 and CLL427) CLL cases by different healthy donor derived CAR T cells including HD-αR110-CAR and anti-TSHR control CAR T cells (HD-αR110-CAR) as indicated and as compared to untransduced cells (UTD). The assay was conducted as in Fig. 2b-d; the 24h time point is shown. (C) Quantification of IFN-γ in co-culture supernatants after 24h incubation of indicated target/effector cell combinations of the assay shown in panel b. (D) Quantification of OCI-Ly1-R110 cytolysis mediated by CLL patient-derived CAR T cells as compared to untransduced cells (UTD). The assay was conducted with two CLL patients serving as T cell donors, one with active CLL (CLL433) and one with CLL in remission (CLL453). The assay was performed as described in Fig. 2b-d; the 24h time point is shown. (E) Workflow for the patient-derived xenograft mouse model for CLL. (F) NSG mice were used to assess in-vivo activity of CLL-αR110-CAR T cells from CLL donor CLL425 with IGLV3-21^R110^-positive CLL. Each mouse was injected i.v. with 0.5 million T cells and 20 million CLL cells collected from patient CLL425. Mice were i.p.-treated 10 days later with 7 million CLL-αR110-CAR T cells (n = 6) or untransduced cells (UTD, n = 6) from the same patient. Mice were then sacrificed at week 3 post CAR T cell injection. Only n = 5 measurements are shown for the CLL-CAR-αR110 T cell treated group since one mouse died of unknown reason. All bar plots represent the indicated mean ± SD. Statistics: one-sided t test.

### Anti-IGLV3-21^R110^ CAR T cells do not mediate B cell toxicity in vitro and in vivo

Finally, we assessed the effect of HD-αR110-CAR T cells on polyclonal healthy B cells in vitro and in vivo. First, we isolated polyclonal B cells from healthy individuals and subjected them to co-culture killing assays with the different CAR T products. While polyclonal B cells were eradicated by HD-αCD19-CAR T cells, HD-αR110-CAR T cells spared this non-malignant compartment demonstrating the epitope-specificity of our targeting approach (Figure 4A). Next, we used two humanized mouse models to show epitope selectivity of HD-αR110-CAR T cells with simultaneous sparing of healthy polyclonal human B cells in vivo (Figure 4B,D). In the first model, human PBMCs were injected intravenously in NSG mice followed by CAR T cell injection seven days later. Quantification of blood circulating human B cells (CD19^+^CD20^+^) showed their persistence after eight days and subtotal eradication of B cells after 13 days when mice were treated with HD-αCD19-CAR T cells (Figure 4C). In contrast, treatment of mice with HD-αR110-CAR T cells did not reduce blood B cell counts after 13 days (Figure 4C). In the second model, NFA2 mice were injected intraperitoneally with human PBMCs and either HD-αCD19-CAR or HD-αR110-CAR T cells (Figure 4D). Quantification of human B cells in peritoneal lavage after 16 hours showed subtotal reduction of B cell counts (CD19^+^) when NFA2 mice were treated with HD-αCD19-CAR T but not with HD-αR110-CAR T cells (Figure 4E).

**Figure 4.**
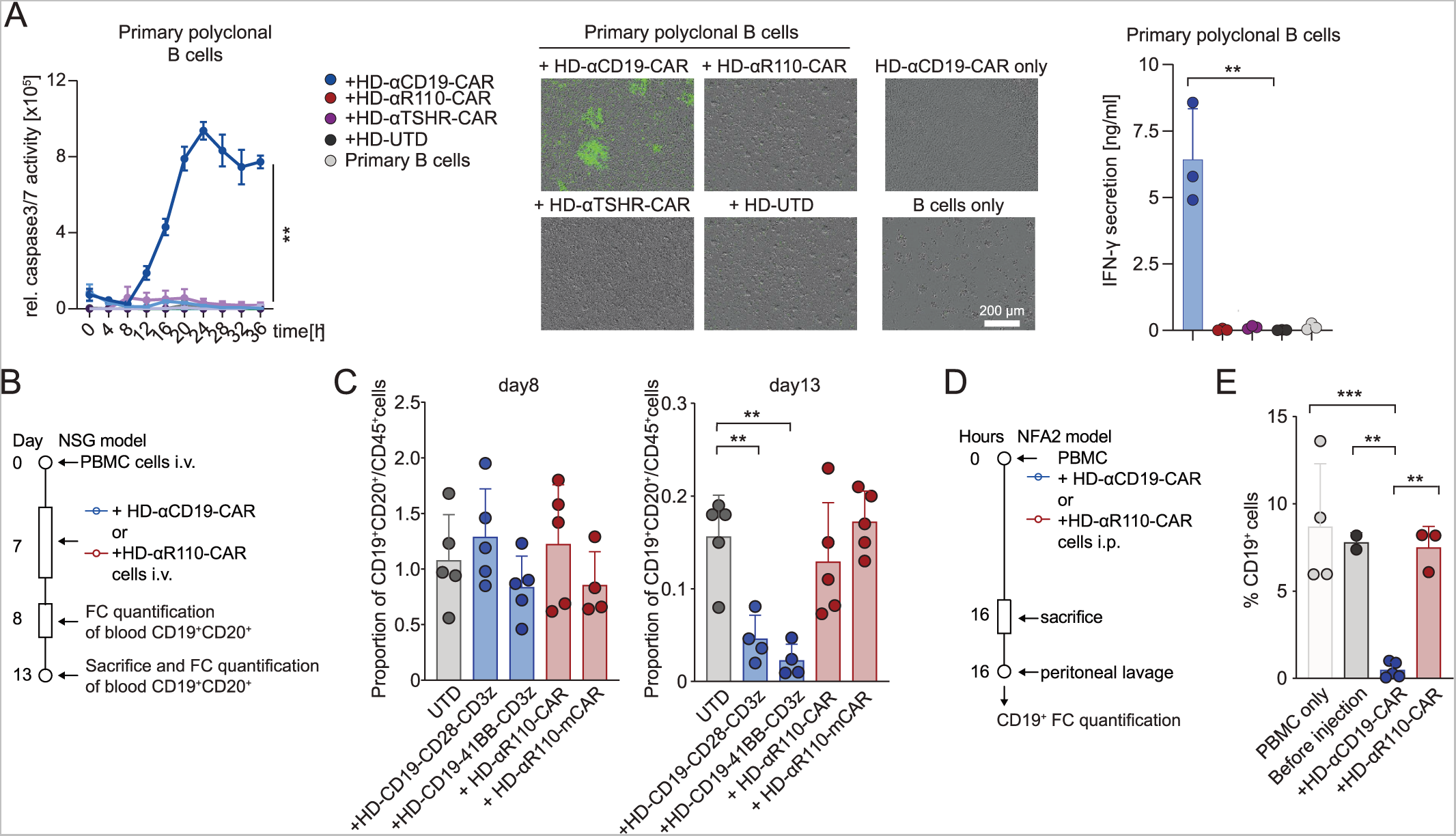
IGLV3-21^R110^ directed CAR T cells spare polyclonal healthy human B cells in vitro and in two humanized mouse models. (A) Quantification of polyclonal B cell cytolysis mediated by healthy donor-derived CAR T cells as compared to untransduced cells (UTD). The assay was conducted as those in Fig. 2b-d. Representative images show CAR T cell-mediated cytolysis after 24h. All conditions indicated have been plotted, negative control conditions overlap. (B)-(E) Humanized mouse models to test the specificity of the αR110-CAR T cells. (B) Workflow of humanized NSG model. (C) Human PBMCs were injected intravenously (i.v.) in NSG mice followed by i.v. injection of UTD, HD-αR110-CAR T, or HD-αCD19-CAR T cells (for all n = 5) on day 7. The abundance of CD19^+^CD20^+^ B cells relative to all CD45^+^ cells was quantified in the blood on day 8 and 13 using flow cytometry (FC). (D) Workflow of humanized NFA2 model. (E) Human PBMCs were intraperitoneally (i.p.) injected into NFA2 mice (n=3-4) along with HD-αR110-CAR T or HD-αCD19-CAR T cells. After 16 h mice were sacrificed, peritoneal cells were harvested by peritoneal lavage and quantified using FC. All bar plots represent the indicated mean ± SD. Statistics: one-sided t test.

## Discussion

CAR T cells are now a mainstay of therapy for B cell malignancies that results in long-term remission in many patients and has the potential of cure^48–53^. The potency of B cell-directed CAR T cell therapy is also highlighted by the recently reported success of treating refractory systemic lupus^54^. Since current CAR T products under development target common (co-) activation markers on the surface of B lymphocytes like CD19, CD20, CD22 and BCMA, these therapies have the drawback of eradicating the B cell lineage or substantial parts of it. As a consequence, B cell-depleted patients are more susceptible for infections complicating clinical management. Another drawback of such strategy is that most B cell-related antigens are not functionally relevant to the cancer cell. As a consequence, target expression, can get lost or mutated to prevent CAR engagement. An approach hitting a disease-driving antigen would stand lower chances of loss or downregulation. Given that target antigen modulation or loss under therapeutic pressure is a well-documented clinical issue affecting up to 50% of patients^58^, the clinical significance of such an approach becomes apparent.

To respond to these challenges, we engineered a selective CAR T construct that targets a recurrent oncogenic point mutation in the BCR light chain of malignant CLL cells. We demonstrate that this approach is feasible and provide in vitro and in vivo data for the selectivity of these CAR T cells towards engineered cell lines as well as primary CLL cells from patients with the IGLV3-21^R110^ mutation. The observed efficacies were comparable to those observed for CD19-directed CAR T cells. Most importantly, we did not observe CAR T-mediated cytotoxicity towards healthy B cells in two humanized mouse models, arguing for the safety of our CAR T product.

Potential clinical applications of IGLV3-21^R110^-targeting CAR T cells range from treatment of relapsed/refractory disease to consolidation after insufficient response to standard first-line CLL treatment. The curative potential of our approach, however, needs further evaluation in clinical trials. While the here applied CLL xenograft model shows nearly complete eradication of engrafted primary CLL R110^+^ B lymphocytes cells under anti-IGLV3-21^R110^ CAR T therapy, primary CLL mice models come with limitations hampering survival analyzes as surrogate for long-term clinical efficacy in CLL patients. The main drawback here - besides the technically challenging requirement of concomitant engraftment and expansion of autologous T cells – is the fact, that CLL cells only transiently engraft and rather survive in a steady state than proliferate in the host^59–62^. To the best of our knowledge any long-term therapeutic model using primary CLL cells in vivo has yet to be found.

In summary, we have developed and provided evidence of the activity and selectivity of what we believe to be the first genuinely tumor-specific, biomarker-driven cellular targeting approach for a hematological malignancy. Our work aligns with the endeavors of various research groups currently striving to create CAR T cells with highly specific targeting for lymphoma cells^55,56^ or autoreactive B cells^57^.

## Acknowledgements

The authors thank Christoph Wosiek, Aline Patzschke and Yiqing Du for excellent technical assistance. Financial support from the Deutsche Forschungsgemeinschaft (DFG BI 1711/4-1 to M.B.) and intramural Roux program funding (to M.B.) is acknowledged. We acknowledge the iFlow Core Facility of the LMU University Hospital and Alexander Navarrete-Santos of the Cell Sorting Core Facility of the Martin-Luther-University (MLU) Halle-Wittenberg for assistance with the generation of flow cytometry data. We also thank Nadine Bley and the Core Facility Imaging of the MLU for help with live cell imaging. This study was further supported by the International Doctoral Program iTarget: Immunotargeting of Cancer funded by the Elite Network of Bavaria (S.K.), the Melanoma Research Alliance Grants 409510 (to S.K.), the Marie-Sklodowska-Curie Program Training Network for Optimizing Adoptive T Cell Therapy of Cancer funded by the H2020 Program of the European Union (Grant 955575, to S.K.), the German Cancer Aid (AvantCAR.de to S.K.), the Else-Kröner-Fresenius-Stiftung (to S.K.), the Wilhelm-Sander-Stiftung (to S.K.), the Deutsche José Carreras Leukämie-Stiftung (to S.K.), the Fritz-Bender-Stiftung (to S.K.) LMU Munich’s Institutional Strategy LMUexcellent within the framework of the German Excellence Initiative (to S.K.), the Bundesministerium für Bildung und Forschung Projects Oncoattract, CONTRACT and Beyondantibody (S.K.), by the Bavarian Research Foundation (BAYCELLATOR to S.K.), by the European Research Council Grant 756017 and 101100460 (to S.K.), Deutsche Forschungsgemeinschaft (DFG; KO5055-2-1 and 510821390 to S.K.) and by the SFB-TRR 338/1 (2021–452881907 to S.K.).

## Author contributions

Idea & design of research project: SK, MB; Supply of critical material (e.g. patient material, mouse models, cohorts): MB, NC, MDM, TN, MH, OC, LE, JM; Establishment of Methods: SK, MB, FM, MA, CS, LP, S-SC, NC, TN, MH, MDvM; Experimental work: FM, MA, CS, S-SC, LP, TZ, SeS, AH, JD, SoS; Analysis and interpretation of primary data: MB, SK, CS, FM, MA, LP, S-SC, NC; Drafting of manuscript: MB, SK, CS. All authors reviewed and revised the manuscript.

## Data availability statement

Sequences of humanized scFVs are subject to ongoing patent application (Application Number: EP22156205.1 / EP22186810.2). NGS data is deposited at the European

Nucleotide Archive (ENA) under the accession number PRJEB65274. All other data supporting the findings of this study are provided with this paper.

## Code availability

No code has been developed for this study.

