## Supplemental Table 1 for "Mutation-specific CAR T cells as precision therapy for IGLV3-21^R110^ expressing high-risk chronic lymphocytic leukemia": Supplemental _material.pdf

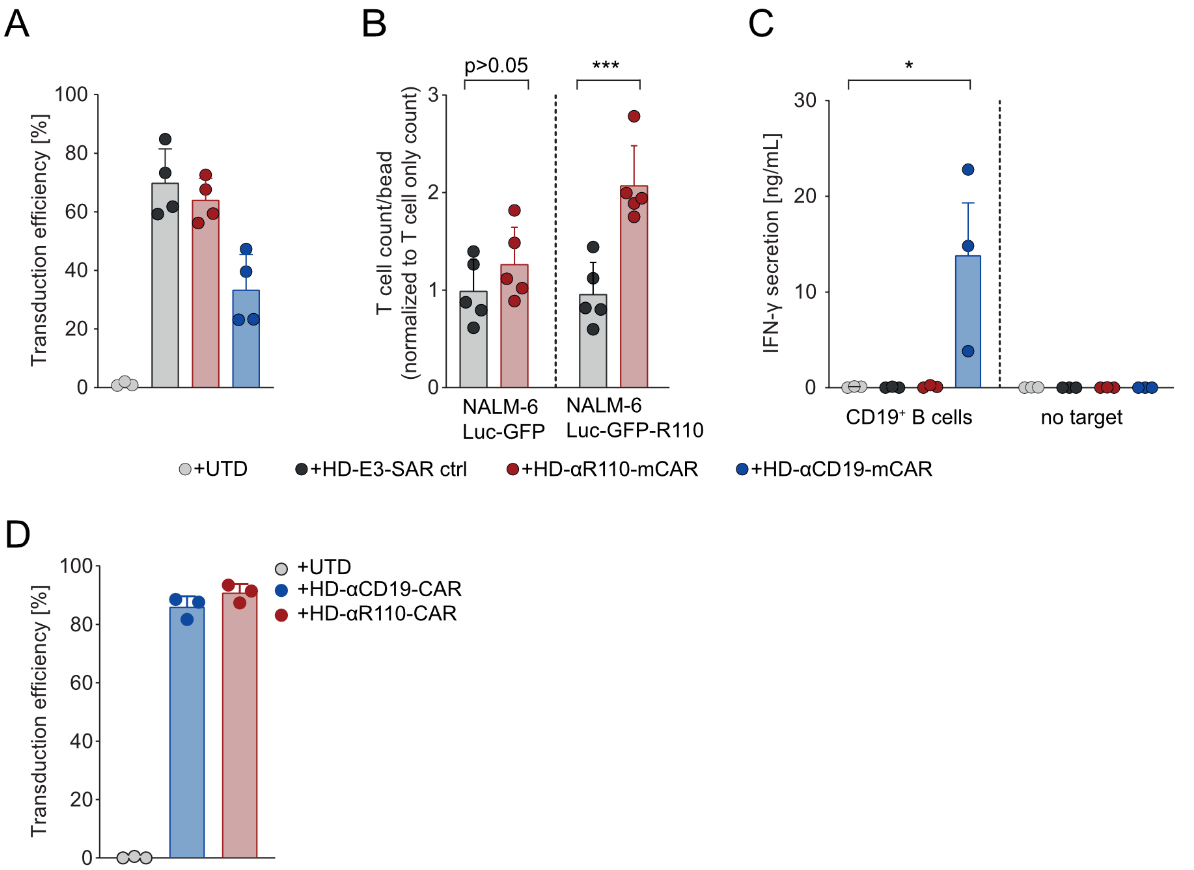


**Supplemental Figure 1.** (A) Efficiency of the murine CAR construct transduction into human T cells. (B) Expansion of HD-αR110-mCAR T cells in co-culture with NALM-6 Luc-R110. Following 48h of co-culture with NALM-6 Luc-R110, $CD3^+$ T cell count per counting bead was assessed by flow cytometry. Counts were normalized to T cells grown as monoculture. (C) Quantification of IFN-γ secretion in cell culture supernatants after 48h co-culture of $CD19^+$ B cells with indicated CAR T cells. (D) Efficiency of the humanized CAR construct transduction into human T cells. All bar plots represent the indicated mean ± SD. Statistics: one-sided t test.
